# Ancient DNA perspectives on North American cervids identify fluctuations in genetic diversity

**DOI:** 10.64898/2026.06.11.731526

**Authors:** Camille Kessler, Oliver Haddrath, Christopher N. Jass, Burton K. Lim, Aaron B.A. Shafer

## Abstract

North American fauna has experienced major changes over the last two million years. Glacial cycles, animal migrations, peopling waves, and, more recently, European colonisation, caused demographic fluctuations, divergence, and wildlife extinctions. Caribou (*Rangifer tarandus*), moose (*Alces alces*), wapiti (*Cervus canadensis*), white-tailed (*Odocoileus virginianus*) and mule deer (*O. hemionus*) are the five extant cervids in Northern America, with unique dispersal patterns and conservation statuses. Here, we used DNA analysis of ancient, historical and modern individuals of these Cervidae to retrace evolutionary processes through time. We show that present-day eastern Canada caribou share ancestry with a 30-thousand-year-old sample from western Canada consistent with the species’ complex dispersal history as suggested by literature. Our results suggest rapid expansion of moose following their arrival on the continent ∼15 thousand years ago. Data on wapiti shows a recent loss of diversity consistent with its large-scale extirpation after European colonisation. Results on *Odocoileus* species are in line with their recognised mito-nuclear discordance. Our results exemplify the utility of chronological sampling to gain a better understanding of how external pressures drive genomic changes.

**Significance statement:** North America has had a dynamic history over the last two million years, with glacial cycles, peopling waves, mammalian extinction events, and European colonisation impacting the regional fauna, including the five deer species currently inhabiting Canada and the USA. It remains unclear how those events impacted each species’ genetic diversity and evolutionary history. Analysis of DNA from samples spanning the last 30 thousand years show signs of expansion after the Last Glacial Maximum for several species, but more recent data shows higher population structure and lower diversity in some cases. Our results indicate each deer species had a unique evolutionary history on the continent, with what appeared to be larger subpopulation ranges during the Holocene whereas more recent times involved more pressures, likely in the form of habitat fragmentation caused by human impact.

## Introduction

Over the last two million years, North America has experienced intense climatic oscillations that caused drastic local and global environmental changes (Allen et al. 2020). The peopling of the continent happened at the very end of these oscillations, around the time of the Last Glacial Maximum (LGM; (Willerslev and Meltzer 2021), and was closely followed by an extensive megafaunal extinction event approximately 11,000 years ago (kya) (Meltzer 2020). The Holocene entailed climatic changes of lower intensity (Shuman and Marsicek 2016), but since the 1500’s, colonisation from Europeans affected wild populations through harvesting, logging, and farming practices (Laliberte and Ripple 2004). Altogether, this dynamic history influenced North American fauna, causing demographic fluctuations, population divergence and range shifts (Lorenzen et al. 2011).

The development of ancient and historical DNA methods allows for sampling of populations before and after an event of interest, and for accurate description of the impact of those events on the genome. Extinct Proboscidea are probably the group that have been most scrutinised with ancient DNA (aDNA), and exemplify the versatility of such data to study admixture, adaptation and selection (e.g.,(van der Valk et al. 2021; van der Valk et al. 2022). Therefore, by using aDNA data, we can retrace the onset of selection, identify drivers of adaptation, and measure changes in genetic diversity through time.

Northern America (Canada & USA) is home to five extant Cervidae species: caribou *(Rangifer tarandus)*, moose *(Alces alces)*, wapiti *(or elk, Cervus canadensis)*, white-tailed deer (WTD; *Odocoileus virginianus)* and mule deer (MD; *O. hemionus)*, each with unique evolutionary histories on the continent. As the range of the North American caribou fluctuated with climatic oscillations, refugial populations from Beringia and south of the ice sheets met and mixed in subsequent dispersal waves (Taylor et al. 2021; Canteri et al. 2025). Moose and wapiti entered the continent after the LGM (Meiri et al. 2020; Mackiewicz et al. 2022) and while information is lacking for wapiti, in moose this was accompanied with a slight genetic bottleneck followed by rapid diversification (Meiri et al. 2020). Wapiti was strongly impacted by human activities after the European colonisation to the point of regional extirpations (O’Gara and Dundas 2002). Both *Odocoileus* sister species originated in North America, and while WTD populations fluctuated over time, MD population size appears to have remained low but stable (Kessler and Shafer 2024). The population size of both species crashed at the turn of the 20^th^ century, and while WTD quickly recovered with minimal genetic loss, there seems to have been no notable recovery in MD (Kessler and Shafer 2024).

Here, we analysed DNA from ancient (> 500 years ago), historical (> 50 years ago) and contemporary samples to explore evolutionary processes for five extant North American Cervidae. Given their specific evolutionary histories, we expect patterns of repeated dispersal waves in caribou, possibly represented by genetic clustering inconsistent with geography. In moose and wapiti, we expected to find signs of rapid expansion, in the form of short phylogenetic branches. Lastly, we predicted a loss in diversity in modern wapiti and MD populations due to the bottlenecks suffered after the European colonisation, whereas the diversity in WTD should have remained stable over time given the species’ large effective population size.

## Results and discussion

North American wildlife evolved through glacial cycles, peopling waves, extinction events, and European colonisation. Here, we used chronological sampling to assess how such events shaped the evolutionary histories of five Cervidae. We analysed ten museum samples dating between 200 and 30k calibrated years before present (cal yBP; **Figure S1, Table S1**). All sequences produced low to medium coverage for mtDNA (1-11x; **Table S2**), whereas nine samples (excluding Odo_ON01) were successfully sequenced at ultra-low coverage at the whole-genome level (0.07–0.9x; **Table S3**), consistent with other aDNA studies (e.g. Skovrind et al. 2024; Arnold et al. 2025), and which presented patterns of DNA damage (**Figure S2, Table S3**).

We successfully extracted DNA from a caribou sample dating approximately 31k cal yBP (Rt_AB03; **Figure S1, Table S1**), which is, to the best of our knowledge, the oldest *Rangifer* analysed with aDNA. Analyses at the nuclear and mitochondrial level consistently showed this Pleistocene sample as closest to modern samples from Ontario, and haplotype network suggests high divergence between all samples (**Figure 1**). This is consistent with the complex dispersal history of North American caribou (Canteri et al. 2025), and points to ancient intermixing of refugial caribou populations.

**Figure 1:**
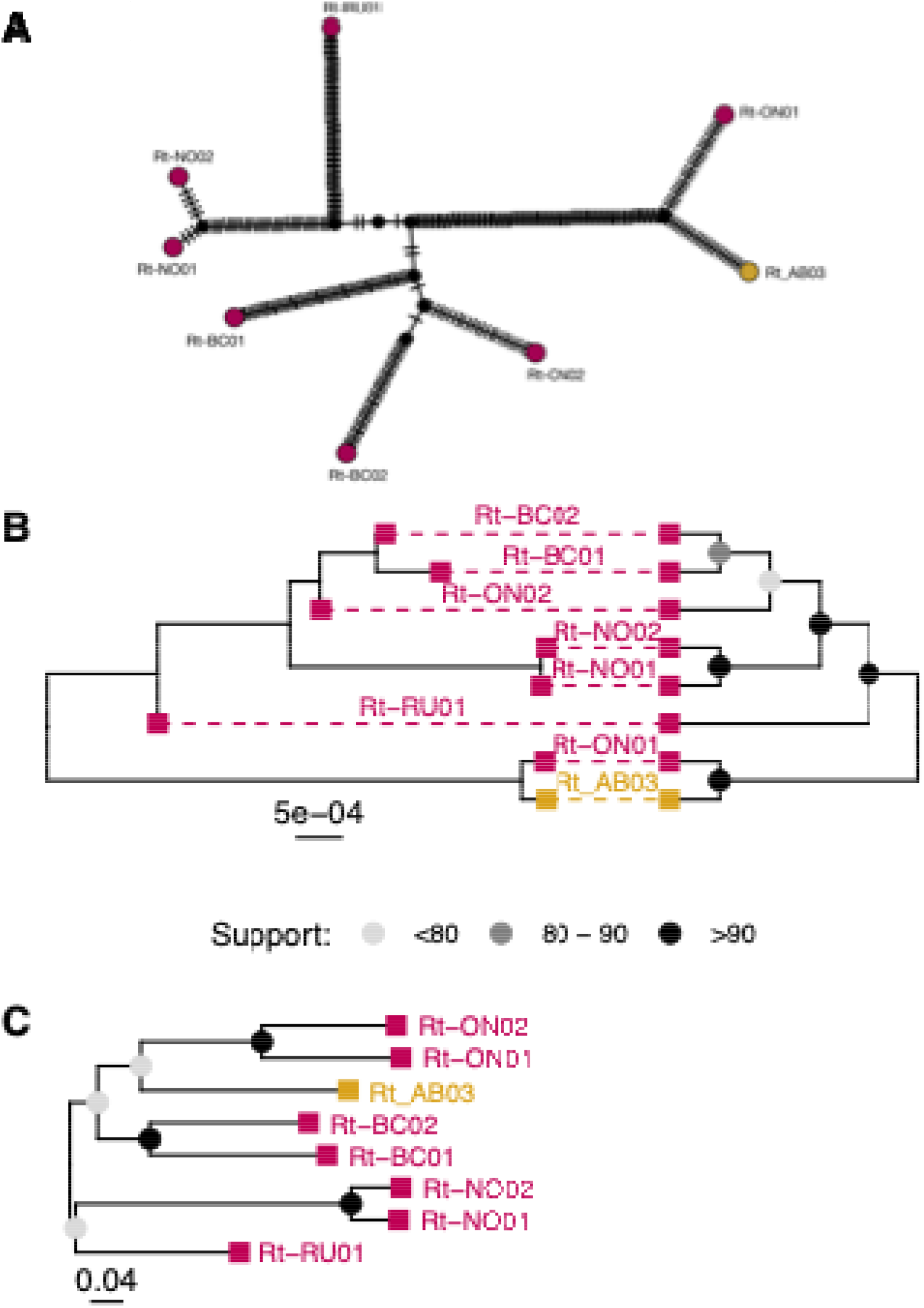
Caribou haplotype network (A), mitochondrial phylogenetic tree and cladogram (B), and nuclear phylogenetic tree (C). Trees in (B) represent the same data in the form of a phylogenetic tree and a cladogram to help understand phylogenetic placement in case of short branches. Node colour indicates support and sample colour denotes modern (burgundy) and museum (mustard) samples.

Our sampling included three ancient wapiti samples (**Figure S1, Table S1**). While all museum samples were labelled *C. elaphus* (**Table S1**), we considered them *C. canadensis* as this species was recognised as distinct from its European counterpart (Pitra et al. 2004). The most ancient (Ce_AB05, ∼12k cal yBP) grouped with modern Wyoming wapiti in the mitochondrial analyses (**Figure 2**), but nuclear analyses show no clear pattern (**Figure S3**). Mitochondrial analyses of Ce_AB07 (∼4000 cal yBP) showed it clustering with modern samples from Minnesota (**Figure 2**), whereas the nuclear phylogeny placed it as basal to the modern Wyoming individuals (**Figure S3**). The most recent sample (Ce_AB02, ∼1000 cal yBP) is closest to European red deer in all phylogenies (**Figure 2B, Figure S3**), whereas it is closest to modern North American samples in the haplotype network (**Figure 2A**). Museum samples presented a higher genetic diversity than modern North American wapiti (**Table S6**), and slightly higher genetic distance between their haplotypes (**Figure 3**). This suggests higher ancestral diversity, consistent with the drastic regional declines of the species in the 19^th^ century (O’Gara and Dundas 2002; Speller et al. 2014).

**Figure 2:**
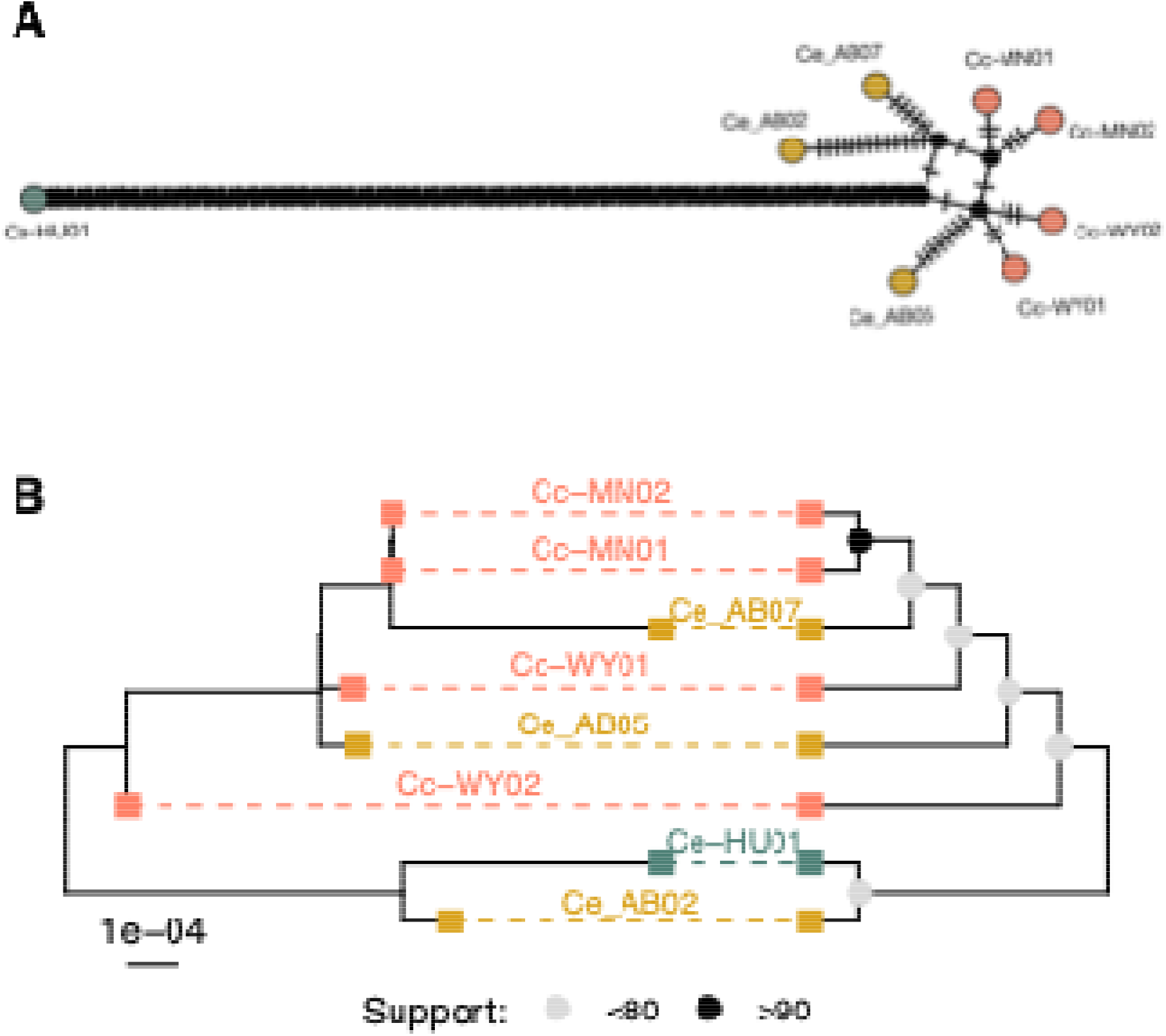
Wapiti mitochondrial analyses. Haplotype network (A) and phylogenetic analyses (B). Both trees represent the same data in the form of a phylogenetic tree and a cladogram to help understand phylogenetic placement in case of short branches. Node colour indicates support and sample colour denotes modern wapiti (orange), modern red deer (dark green) and museum (mustard) samples.

**Figure 3:**
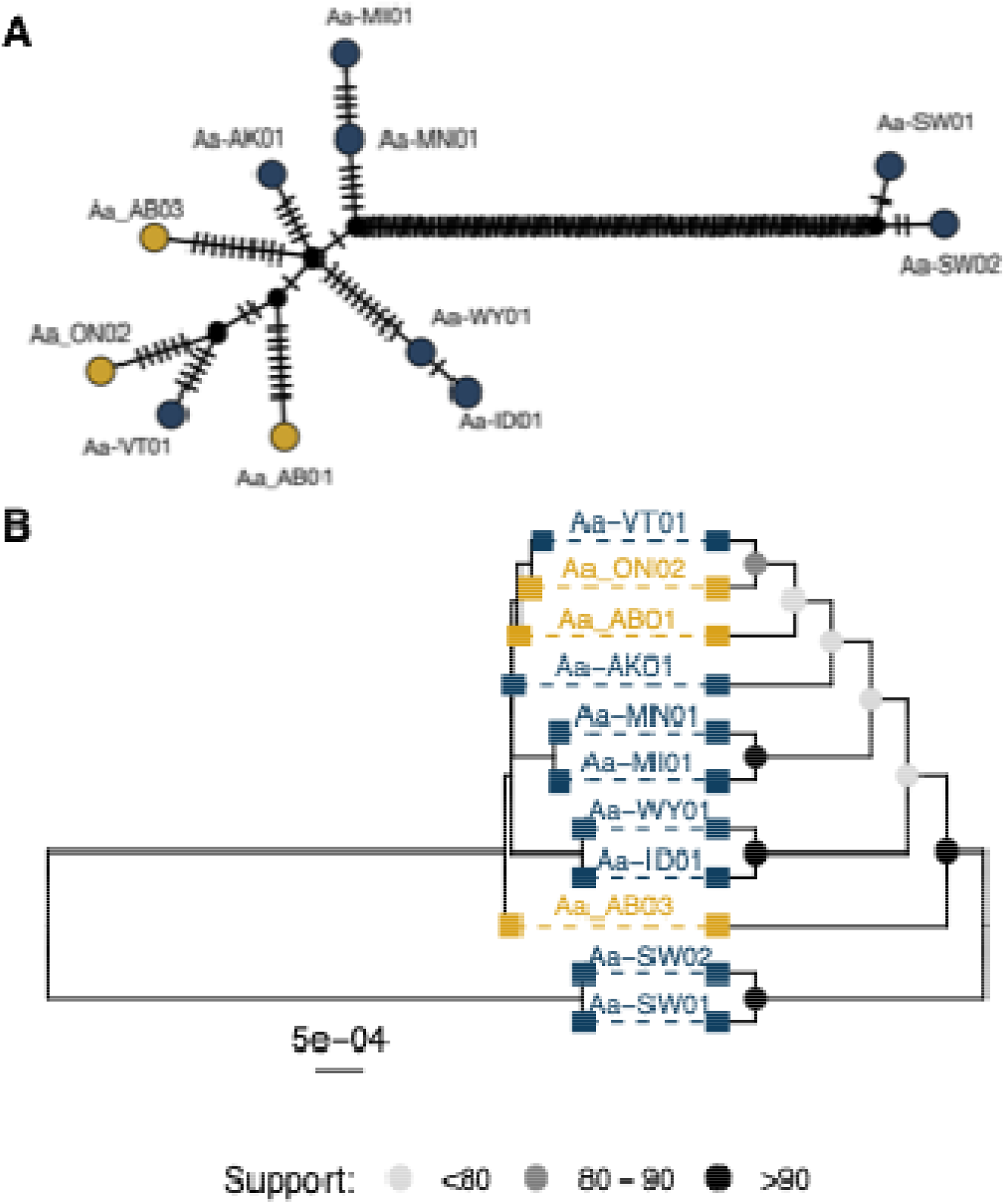
Moose mitochondrial analyses. Haplotype network (A) and phylogenetic analyses (B). Both trees represent the same data in the form of a phylogenetic tree and a cladogram to help understand phylogenetic placement in case of short branches. Node colour indicates support and sample colour denotes modern (dark blue) and museum (mustard) samples.

We sampled two historical and one ancient moose (**Figure S1, Table S1**). The ancient sample Aa_AB01 (∼11,500 cal yBP) represents the oldest, directly dated moose from Alberta, it is basal to the modern Vermont and the historical Ontario samples in the mitochondrial analyses (**Figure 3**). The historical sample Aa_AB03 (∼400 cal yBP) is placed as basal to all North American individuals in the mitochondrial phylogeny, and is nested within the North American clade on the haplotype network (**Figure 3**). Lastly, the Ontario historical sample (Aa_ON02, ∼200 cal yBP) is closest to the modern Vermont moose in both mitochondrial analyses (**Figure 3**). The very short branches on the mitochondrial phylogeny supports rapid and recent diversification (**Figure 3**) consistent with previous literature (Meiri et al. 2020). Nuclear phylogenies showed a similar topology for all museum samples that group with Alaska, Idaho and Vermont individuals (**Figure S4**). This consistency despite the disparity in age between the Alberta samples suggests a lack of differentiation for moose populations throughout the Holocene consistent with the observed homogenous structure over time and space in other taxa of the region (Karpinski et al. 2020; Murchie et al. 2022). Finally, museum samples harboured the lowest genetic diversity in moose (**Table S4**).

We successfully sequenced one historical and one ancient *Odocoileus*, one of which was unidentified to the species-level (**Table S1**). The sample Odo_AB02 (∼4000 cal yBP) is found in a clade containing both WTD and MD in the mitochondrial analyses, and within MD at the nuclear level (**Figure 4, Figure S5**). The second sample (Odo_AB05, ∼140 cal yBP) was consistently placed with modern MD individuals. The WTD historical sample (Odo_ON01) was degraded and excluded from nuclear analyses. Its mitochondrial sequence had just over 60% missing data (**Table S2**) and is placed as sister to a modern WTD from Ontario in the mitochondrial phylogenetic analysis (**Figure 4**). Our mitochondrial analyses show no species clustering in the *Odocoileus* genus (**Figure 4, Figure S5**) consistent with those species’ recognised mito-nuclear phylogenetic discordance likely caused by incomplete lineage sorting after a rapid radiation (Cronin 1991; Klicka et al. 2023; Kessler et al. 2025). Genetic diversity in *Odocoileus* was lowest in the MD museum samples and highest in modern WTD (π = 0.09 & 0.15, respectively, **Table S4**). While we had predicted a drop in diversity in modern MD samples caused by a bottleneck about ∼120 years ago (Kessler and Shafer 2024), secondary contact and modern hybridisation with WTD could have buffered diversity, although hybridisation rates in the wild are low (Russell et al. 2021). It could also suggest a sampling bias as both museum samples come from Alberta, where the MD populations might have lower diversity than the species-wide average.

**Figure 4:**
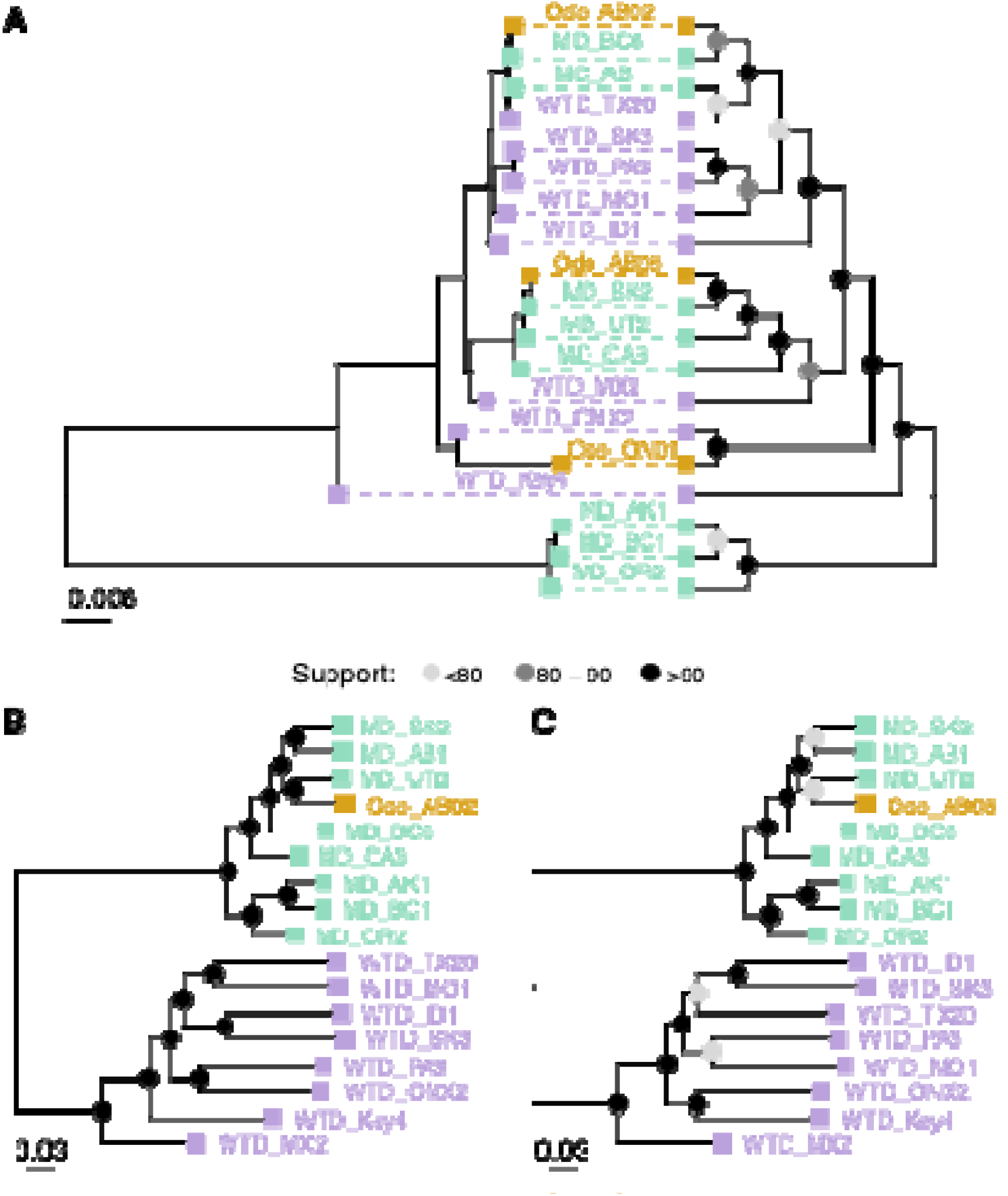
*Odocoileus* phylogenetic analyses at the mitochondrial (A) and nuclear (B-C) levels. Trees in (A) represent the same data in the form of a phylogenetic tree and a cladogram to help understand phylogenetic placement in case of short branches. Node colour indicates support and sample colour denotes represents modern white-tailed deer (violet), modern mule deer (green) and museum (mustard) samples.

In summary, we used museum and modern samples to identify phylogenetic affinities while also comparing diversity metrics, with modern samples capturing changes that occurred post European colonisation. Using chronological sampling, we were able to retrace the changes in genetic diversity through time for five extant cervids, finding patterns generally confirming existing literature based on retrospective analyses. We showed the utility of chronological sampling to better understand how external pressures drive genomic changes in the wild.

## Acknowledgements

We sampled ancient deer from across Canada, a land which remains a home to many First Peoples, who share an ancestral connection with these species. Trent University is located on the territory of the Michi Saagiig Anishnaabeg; the ROM is located on the territory of the Michi Saagiig Anishnaabeg, the Wendat, the Haudenosaunee Confederacy, and the Anishinaabeg Nation, including the Mississaugas of the Credit First Nation; the RAM is located on and houses objects from the traditional territory of the Cree, Saulteaux, Blackfoot, Métis, Dene and Nakota Sioux. We are grateful to have had the opportunity to live and work on this land, and to benefit from it. We would like to show our respect to the First Peoples and thank them for their care, stewardship, and teachings.

We would like to thank Marianne Dehasque for sharing their laboratory methods and protocols, and Marie-Laurence Cossette for their help in the lab.

This work was supported by ComputeCanada (RRG gme-665-ab); NSERC (RGPIN-2017-03934) and the Canadian Foundation for Innovation (#36905). Camille Kessler was supported by an International Graduate Scholarship and an Ontario Graduate Scholarship.

## Material and methods

### Museum sampling and laboratory procedures

We sampled ten North American museum specimens (**Table S1, Figure S1**). For radiocarbon dating, bone samples were first sent to the Trent Environmental Archaeology Lab for collagen extraction and purification, then to the W. M. Keck Carbon Cycle AMS facility at the University of California, Irvine for radiocarbon dating. We calibrated the resulting C^14^ dates in OxCal 4.4 (Ramsey 2009) using the IntCal20 calibration curve (Reimer 2020).

We performed laboratory procedures in a dedicated aDNA facility at the Royal Ontario Museum (Canada). Bone fragments were first UV-irradiated, then powdered in liquid nitrogen using a mortar and pestle to obtain approximately 50 mg of bone powder. For DNA extraction, we followed the silica column protocol and double-stranded library preparation with USER enzyme treatment from (Dehasque et al. 2022). Then, double-indexing and amplification was performed as in (Díez-del-Molino et al. 2023) . We ran four independent PCR amplifications as follows: 95°C for 2 min, then 12 cycles of 95°C for 15 s, 60°C for 30 s and 68 °C for 1 min and included one extraction and one library prep negative control (ddH_2_O) per batch. Libraries were sequenced for 150 bp paired-end reads on an Illumina Novaseq 6000 with SP flowcells.

### Data processing

For the museum data, we trimmed and merged the raw reads in SeqPrep v1.2 (John 2011) with default settings and a minor source code modification (Palkopoulou et al. 2015), then mapped with bwa aln v0.7.17 (Li and Durbin 2009) and settings optimised for aDNA (-o 2, -n 0.01, -l 16500; (Schubert et al. 2012; van der Valk et al. 2022). Samples were mapped to their respective mitochondrial and whole genome reference assembly (or those of closely related species, **Table S5**). Using SAMtools v1.17 (Li et al. 2009), we excluded reads < 30bp and with a mapping quality < 25, and removed PCR duplicates with samremovedup.py (Palkopoulou et al. 2018). Finally, we assessed quality with bamtools stats v2.5.1 (Barnett et al. 2011), computed final coverage in ANGSD v0.939 (Korneliussen et al. 2014), and identified CpG damage patterns (*--platypus*) with PMDtools v2 (Skoglund et al. 2014). We sexed all samples using autosome/X chromosome read ratio for the species with an chromosome-level reference, and SATC (Nursyifa et al. 2022) for species without.

For modern data, we gathered contemporary resequencing data from 36 published genomes (**Table S6**), which we standardised by selecting 100 million random read pairs per sample using seqtk sample v1.3 (Li 2018). Reads were trimmed, aligned and sorted with Trimmomatic v0.36 (Bolger et al. 2014), bwa-mem v0.7.17 (Li and Durbin 2009) and SAMtools v1.10 (Li et al. 2009), respectively. We excluded duplicated reads using Picard MarkDuplicate v2.23.2 (Broad Institute 2019) and Sambamba view v0.7.0 (Tarasov et al. 2015), and lastly performed local realignment in GATK RealignerTargetCreator and IndelRealigner v4.1.7.0 (McKenna et al. 2010).

### Nuclear DNA phylogeny and genetic diversity

We produced random read sampling data in ANGSD (-doIBS 1), this method calls bases from randomly sampled reads at each position, which controls for the uncertainty and low coverage of the museum data. We called SNPs (*-SNP_pval 1e-6*) filtered for “good”, uniquely mapping reads (-*remove_bads 1 -uniqueOnly 1*), with base quality > 30 and mapping quality > 25 (*-minMapQ 25 -minQ 30*), allowing no missing data *(- minInd $N -maxMis 0)*. To minimise data loss from missingness and low SNPs overlap among museum samples, we generated these data independently for each museum sample together, with all modern individuals.

To generate phylogenies, we first further excluded all singletons, then ran IQ-TREE v1.6.12 (Nguyen et al. 2015; Kalyaanamoorthy et al. 2017; Hoang et al. 2018) with best model selection *(-m MFP)*, SH-like approximate likelihood ratio test *(-alrt 1000)*, ultrafast bootstrap *(-bb 1000)* and hill-climbing nearest neighbour interchange search *(-bnni)*. Consensus trees were plotted using the ggtree R package (R Core Team 2021; Xu et al. 2022).

Lastly, to compute genetic diversity, we performed a second ANGSD call (same filtering as above), including all individuals of a same species group which resulted in fewer SNPs given the low sites overlap in museum samples. Genetic diversity was computed in the ape and pegas R packages (Paradis 2010; Paradis and Schliep 2019), splitting the samples into relevant groups (e.g. Museum, North America, Europe).

### Haplotype networks and phylogenetic analysis of mitochondrial DNA

For mitochondrial DNA (mtDNA) analysis, we produced consensus sequences independently for each sample in ANGSD, selecting the base with highest effective depth and setting a minimum depth of 3 *(-doFasta 3 -setMinDepth 3)*. Consensus sequences from all samples of a same species were aligned in MEGA v11.0.13 (Stecher et al. 2020) using the ClustalW algorithm with default settings. We inferred median joining networks in popART (Leigh and Bryant 2015) with and without samples having over 50% missing data. Finally, to produce a phylogenetic tree, we first removed all gaps and singletons from the alignments and ran IQ-TREE as above.

### Data availability statement

Raw reads for the ancient samples have been deposited on the NCBI Sequence Read Archive as BioProject PRJNA1474373 (reviewer link: https://dataview.ncbi.nlm.nih.gov/object/PRJNA1474373?reviewer=geh4s6iqspt7s9a87lpbjc7qe9). This paper also analyses publicly available data, accession numbers can be found in **Table S3**. All scripts are publicly available on https://gitlab.com/WiDGeT_TrentU/graduate_theses/-/tree/master/Kessler/CH_04

