## Supplementary figures and tables for "Ancient DNA perspectives on North American cervids identify fluctuations in genetic diversity"

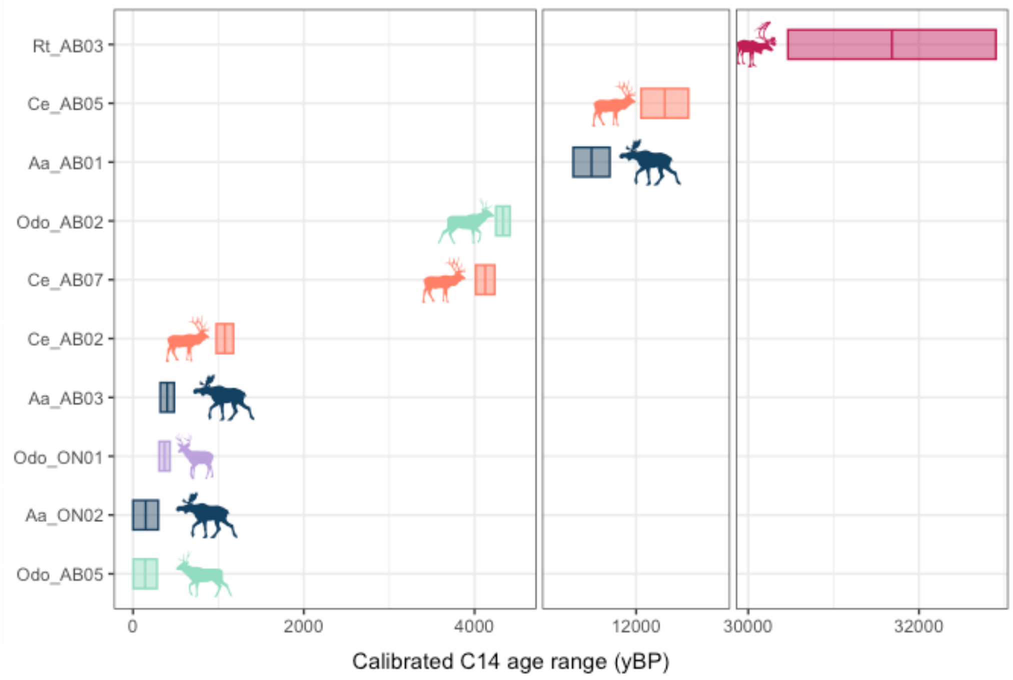


Figure S1: Museum samples calibrated radiocarbon ranges timeline. Coloured by species: caribou in burgundy, wapiti in orange, moose in dark blue, mule deer in green and white-tailed deer in violet.

Figure S2: Damage patterns inferred in PMDtools in each museum sample when mapped to their target species. Deamination at CpG sites identified by PDMtools is unaffected by USER enzyme treatment.

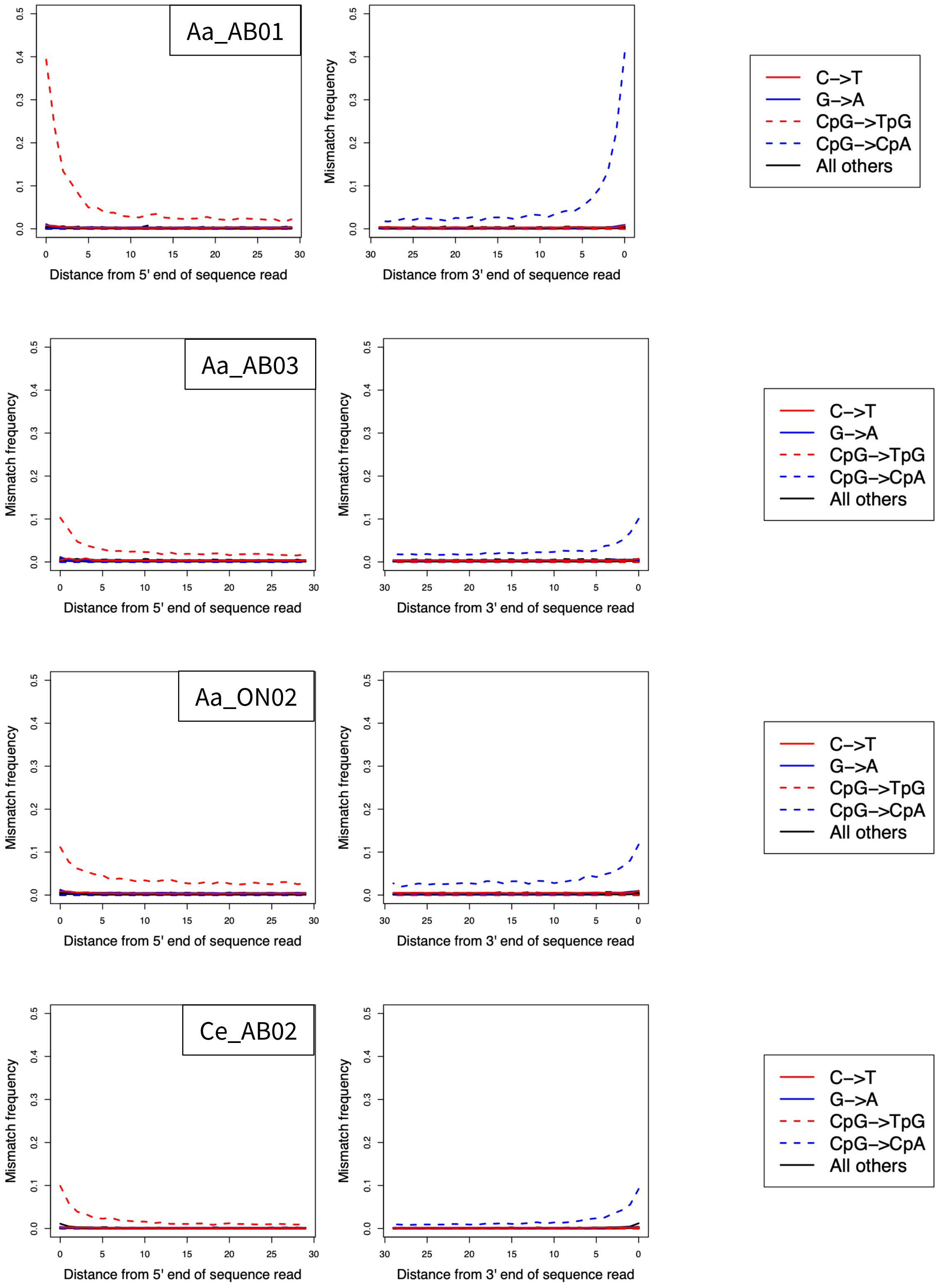

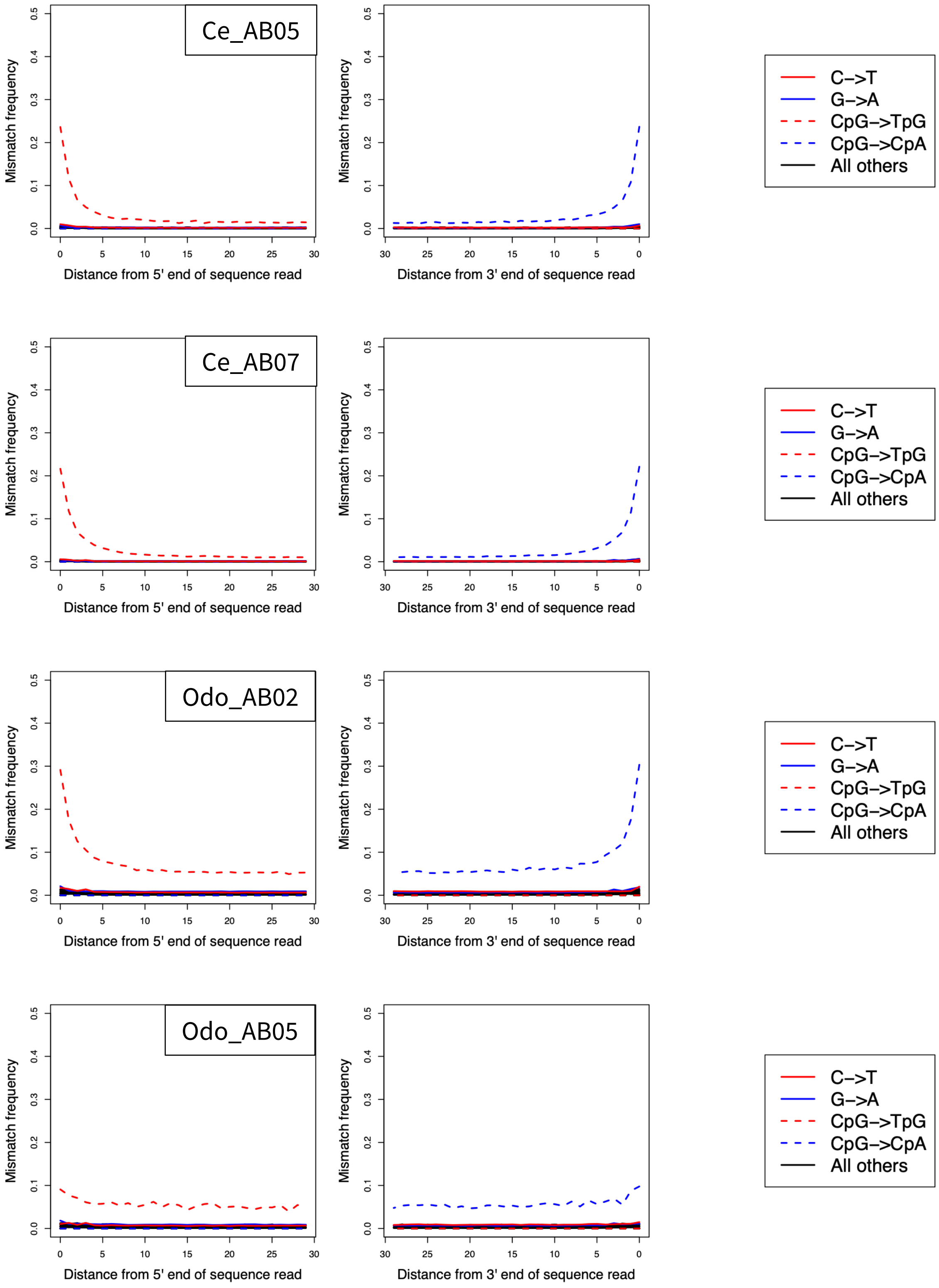
Ce_AB05
Ce_AB07
Odo_AB02
Odo_AB05
Mismatch frequency
0.0
0.1
0.2
0.3
0.4
0.5
0
5
10
15
20
25
30
Distance from 5' end of sequence read
Distance from 3' end of sequence read
C−>T
G−>A
CpG−>TpG
CpG−>CpA
All others

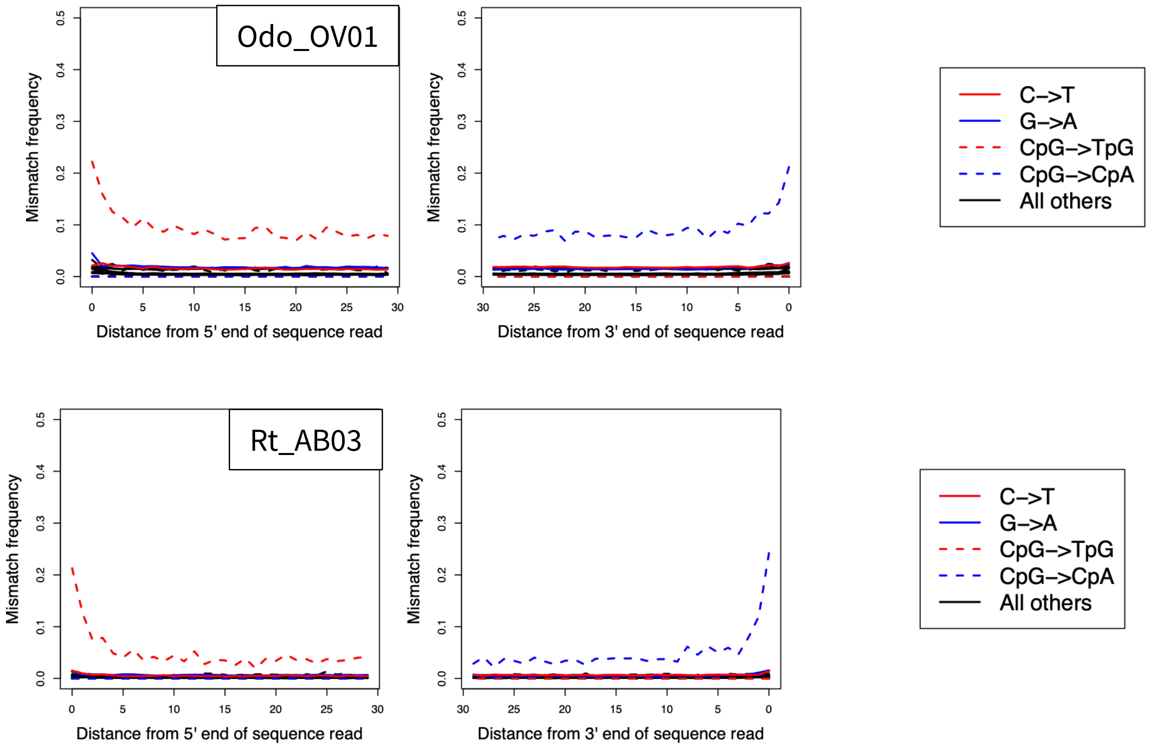
Odo_OV01
Mismatch frequency
Distance from 5' end of sequence read
Distance from 3' end of sequence read
C–>T
G–>A
CpG–>TpG
CpG–>CpA
All others
Rt_AB03
Mismatch frequency
Distance from 5' end of sequence read
Distance from 3' end of sequence read
C–>T
G–>A
CpG–>TpG
CpG–>CpA
All others

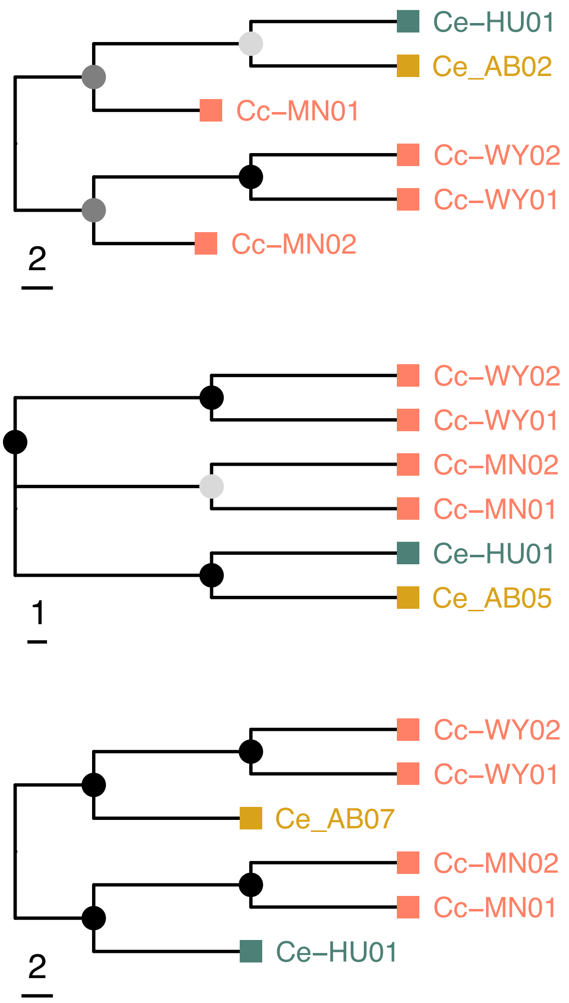


Figure S3: Wapiti nuclear analyses. Per individual phylogenetic tree (left) based on random read sampling. Phylogenetic tree shows node support with coloured node points. Sample colour represents modern wapiti (orange), modern red deer (dark green) and museum (mustard) samples.

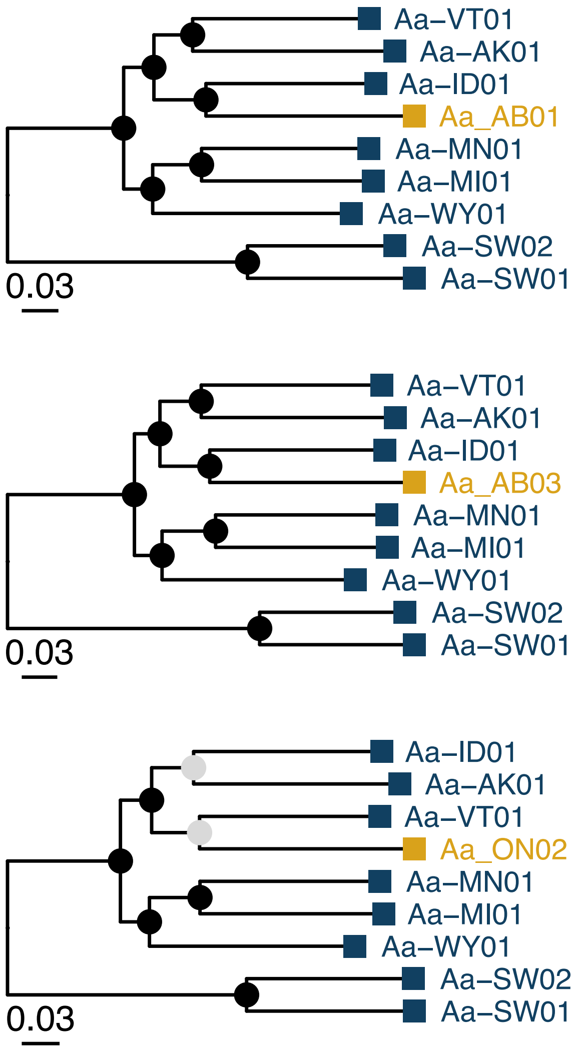


Figure S4: Moose nuclear analyses. Per individual phylogenetic tree (left) based on random read sampling. Phylogenetic tree shows node support with coloured node points. Sample colour represents modern (dark blue) and museum (mustard) samples.

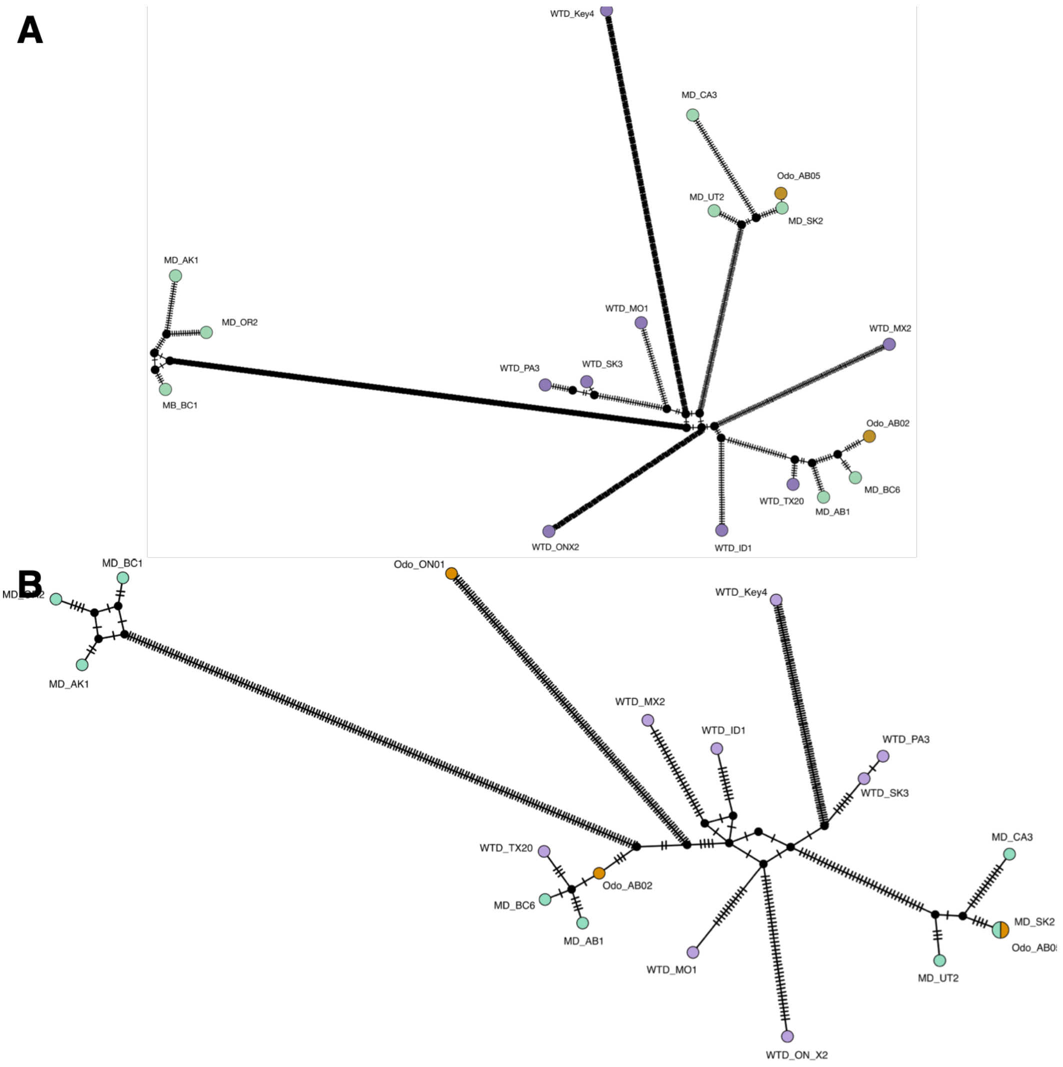


Figure S5: Odocoileus haplotype networks without (A) and with (B) sample with >50% missing data (Odo_ON01). Sample colour represents modern white-tailed deer (violet), modern mule deer (green) and museum (mustard) samples.

Table S1: Museum sample information. Short ID was used in paper and figures for clarity as it contains the species and broad location; specimen ID as used by the museums; provider abbreviation : ROM = Royal Ontario Museum and RAM = Royal Alberta Museum; Species as labelled in museum records; C14 age (BP) raw estimate from KCCAMS labs; Calibrated C14 age as computed in OxCal 4.4.

| Short ID | Specimen ID | Sample source | Provider | Species | C14 age (BP) | Calibrated C14 age range BP (95.4%) | Mean calibrated C14 age BP |
|---|---|---|---|---|---|---|---|
| Aa_AB01 | P19.385.6 | Right mandible | RAM | *Alces alces* | 9,980 ± 45 | 11,263 - 11,692 | 11477.5 |
| Aa_AB03 | P02.9.34 | Left metacarpal | RAM | *Alces alces* | 355 ± 15 | 319 - 482 | 400.5 |
| Aa_ON02 | ROMM33679 | Upper jaw | ROM | *Alces alces* | 205 ± 15 | 296 - out of range | NA |
| Ce_AB02 | P02.9.67 | Left tibia | RAM | *Cervus canadensis* | 1,150 ± 15 | 975 - 1,175 | 1075 |
| Ce_AB05 | P85.10.1 | Skull | RAM | *Cervus canadensis* | 10,435 ± 50 | 12,058 - 12,612 | 12335 |
| Ce_AB07 | P18.236.4 | Partial mandible | RAM | *Cervus canadensis* | 3,770 ± 20 | 4,013 - 4,235 | 4124 |
| Odo_AB02 | P14.9.3 | Right metatarsal | RAM | *Odocoileus hemionus* | 3,905 ± 20 | 4,250 - 4,415 | 4332.5 |
| Odo_AB05 | P16.12.1 | Right mandible | RAM | *Odocoileus sp. indet.* | 155 ± 15 | 4 - 281 | 142.5 |
| Odo_ON01 | ROMM11048 | Mandible | ROM | *Odocoileus virginianus* | 305 ± 15 | 306-433 | 369.5 |
| Rt_AB03 | P94.1.297 | Left metatarsal | RAM | *Rangifer tarandus* | 27,330 ± 470 | 30,465 - 32,901 | 31683 |

Table S2: Ancient samples mitochondrial genome QC; filtering QC refers to final read numbers using minQ > 25, Length >30, no duplicates

| Sample | Mapping | | Filtering | Coverage | |
|---|---|---|---|---|---|
| | Reference species | Mapped reads to self (#) | Fitlered reads (#) | depth | %N in consensus sequence |
| Aa_AB01 | *Alces alces* | 1159 | 1055 | 4.5421 | 1.105 |
| Aa_AB03 | *Alces alces* | 3429 | 2432 | 11.5671 | 0.257 |
| Aa_ON02 | *Alces alces* | 1466 | 1074 | 4.7762 | 2.120 |
| Ce_AB02 | *Cervus canadensis* | 228 | 216 | 1.5861 | 43.982 |
| Ce_AB05 | *Cervus canadensis* | 1027 | 737 | 3.1776 | 5.980 |
| Ce_AB07 | *Cervus canadensis* | 1719 | 1298 | 5.0664 | 1.270 |
| Odo_AB02 | *Odocoileus virginianus* | 1310 | 1138 | 5.2301 | 1.606 |
| Odo_AB05 | *Odocoileus virginianus* | 1312 | 1209 | 6.0828 | 1.041 |
| Ov_ON01 | *Odocoileus virginianus* | 136 | 116 | 1.3198 | 65.717 |
| Rt_AB03 | *Rangifer tarandus* | 1795 | 1304 | 6.0782 | 1.450 |

Table S3: Ancient samples nuclear genome QC; filtering QC refers to final read numbers using minQ > 25, Length >30, no duplicates; IBS parsimony informative sites refers to the number of non-singleton random bases generated for the IQtree analysis

| Sample | Mapping | | | Filtering | | | IBS | Sex ID | |
|---|---|---|---|---|---|---|---|---|---|
| | Reference species | Mapped reads to self (#) | Mapped reads to self (%) | Fitlered reads (#) | Filtered read length (avg) | Depth | Parsimony informative sites | X/A ratio | Inferred sex |
| Aa_AB01 | *Alces alces* | 424790 | 11.5667 | 216876 | 58 | 0.1546 | 21259 | 0.519 | Male |
| Aa_AB03 | *Alces alces* | 1283762 | 4.15461 | 556103 | 72 | 0.4878 | 55708 | 0.515 | Male |
| Aa_ON02 | *Alces alces* | 713289 | 7.19809 | 298525 | 64 | 0.2338 | 31930 | 0.505 | Male |
| Ce_AB02 | *Cervus canadensis* | 1205635 | 60.2798 | 603807 | 66 | 0.0963 | 45837 | 0.993 | Female |
| Ce_AB05 | *Cervus canadensis* | 1382086 | 24.7406 | 486523 | 59 | 0.0696 | 33573 | 0.588 | Male |
| Ce_AB07 | *Cervus canadensis* | 12311879 | 31.844 | 5516610 | 67 | 0.8911 | 369415 | 0.949 | Female |
| Odo_AB02 | *Odocoileus virginianus* | 1502949 | 10.7382 | 828120 | 74 | 0.4225 | 629878 | 0.441 | Male |
| Odo_AB05 | *Odocoileus virginianus* | 303780 | 10.3202 | 181249 | 84 | 0.1046 | 160853 | 0.456 | Male |
| Rt_AB03 | *Rangifer tarandus* | 105255 | 1.2914 | 56501 | 69 | 0.1623 | 11645 | 0.846 | Female |

Table S4: Genetic diversity estimates

| Species | Population | Genetic diversity |
|---|---|---|
| Moose | Museum | 0.16213 |
| | Europe | 0.20467 |
| | North America | 0.20539 |
| Wapiti | Museum | 0.19648 |
| | Europe | NA |
| | North America | 0.17573 |
| *Odocoileus* | Museum | 0.08962 |
| | WTD | 0.14730 |
| | MD | 0.12220 |
| Caribou | Museum | NA |
| | Eurasia | 0.23470 |
| | North America | 0.31995 |

Table S5: Reference genome information per species.

| Reference species | Samples mapped | Complete reference | Mitochondrial reference |
|---|---|---|---|
| **Alces alces** | Moose | GCA_015832495.2 | NC_020677.1 |
| **Cervus canadensis** | Wapiti and red deer | GCF_019320065.1 | NC_050863.1 |
| **Rangifer tarandus** | Caribou | GCA_949782905.1 | NC_015247.1 |
| **Odocoileus virginianus** | Mule deer & white-tailed deer | GCA_014726795.1 | OX460346.1 |

Table S6: Modern sample information as retrieved from NCBI SRA metadata information. Sample ID designated for clarity.

| Species | Sample ID | Accession | Locality | Sex |
|---|---|---|---|---|
| Alces alces ● | Aa-AK01 | SRR6079187 | Alaska | Male |
| Alces alces ● | Aa-ID01 | SRR6079177 | Idaho | Male |
| Alces alces ● | Aa-MI01 | SRR18899225 | Michigan | NA |
| Alces alces ● | Aa-MN01 | SRR18899222 | Minnesota | NA |
| Alces alces ● | Aa-SW01 | ERR12087977 | Sweden | NA |
| Alces alces ● | Aa-SW02 | ERR12087904 | Sweden | NA |
| Alces alces ● | Aa-VT01 | SRR6079181 | Vermont | Male |
| Alces alces ● | Aa-WY01 | SRR6079200 | Wyoming | Female |
| Cervus canadensis ● | Cc-WY01 | SRR12450513 | Wyoming | Female |
| Cervus canadensis ● | Cc-MN01 | SRR12450505 | Minnesota | Male |
| Cervus canadensis ● | Cc-MN02 | SRR12450502 | Minnesota | Female |
| Cervus canadensis ● | Cc-WY02 | SRR12450506 | Wyoming | Female |
| Cervus elaphus ● | Ce-HU01 | SRR956941 | Hungary | Male |
| Rangifer tarandus ● | Rt-BC01 | SRR27590283 | British Columbia | NA |
| Rangifer tarandus ● | Rt-BC02 | ERR11471728 | British Columbia | NA |
| Rangifer tarandus ● | Rt-ON01 | ERR11471702 | Ontario | Male |
| Rangifer tarandus ● | Rt-ON02 | SRR24951207 | Ontario | NA |
| Rangifer tarandus ● | Rt-NO01 | SRR24951210 | Norway | NA |
| Rangifer tarandus ● | Rt-NO02 | SRR15459420 | Norway | NA |
| Rangifer tarandus ● | Rt-RU01 | SRR15459424 | Russia | NA |
| Odocoileus hemionus ● | Oh_AB1 | SRS12707053 | Alberta | Female |
| Odocoileus hemionus ● | Oh_AK1 | SRS12707054 | Alaska | NA |
| Odocoileus hemionus ● | Oh_BC1 | SRS12707032 | British Columbia | Female |
| Odocoileus hemionus ● | Oh_BC6 | SRS12707055 | British Columbia | Male |
| Odocoileus hemionus ● | Oh_CA3 | SRS12707057 | California | NA |
| Odocoileus hemionus ● | Oh_OR2 | SRS12707035 | Oregon | NA |
| Odocoileus hemionus ● | Oh_SK2 | SRS16492111 | Saskatchewan | NA |
| Odocoileus hemionus ● | Oh_UT2 | SRS12707038 | Utah | Male |
| Odocoileus virginianus ● | Ov_ID1 | SRS16492129 | Idaho | Female |
| Odocoileus virginianus ● | Ov_Key4 | SRS16492093 | Florida | NA |
| Odocoileus virginianus ● | Ov_MO1 | SRS16492102 | Missouri | Male |
| Odocoileus virginianus ● | Ov_MX2 | SRS12707043 | Mexico | Male |
| Odocoileus virginianus ● | Ov_ONX2 | SRS16492116 | Ontario | Female |
| Odocoileus virginianus ● | Ov_PA3 | SRS12707047 | Pennsylvania | Female |
| Odocoileus virginianus ● | Ov_SK3 | SRS12707051 | Saskatchewan | Male |
| Odocoileus virginianus ● | Ov_TX20 | SRS16492126 | Texas | Female |
